# Fibroblast growth factor 9 stimulates neurite outgrowth through NF-kB signaling in striatal cell Huntington’s disease models

**DOI:** 10.1101/2020.07.17.207837

**Authors:** Issa Olakunle Yusuf, Hsiu-Mei Chen, Pei-Hsun Cheng, Chih-Yi Chang, Shaw-Jenq Tsai, Jih-Ing Chuang, Chia-Ching Wu, Bu-Miin Huang, H. Sunny Sun, Chuan-Mu Chen, Shang-Hsun Yang

**Affiliations:** Taiwan International Graduate Program in Interdisciplinary Neuroscience, National Cheng Kung University and Academia Sinica, Taipei 11529, Taiwan; Institute of Clinical Medicine, National Cheng Kung University, Tainan 70101, Taiwan; Department of Physiology, College of Medicine, National Cheng Kung University, Tainan 70101, Taiwan; Institute of Basic Medical Sciences, National Cheng Kung University, Tainan 70101, Taiwan; Department of Cell Biology and Anatomy, National Cheng Kung University, Tainan 70101, Taiwan; Institute of Molecular Medicine, National Cheng Kung University, Tainan 70101, Taiwan; Department of Life Sciences, College of Life Sciences, National Chung Hsing University, Taichung 40227, Taiwan

**Keywords:** Fibroblast Growth Factor 9, NF-κB signaling, neurite outgrowth, Huntington’s disease, striatal cells

## Abstract

Proper development of neuronal cells is important for brain functions, and impairment of neuronal development may lead to neuronal disorders, implying that improvement in neuronal development may be a therapeutic direction for these diseases. Huntington’s disease (HD) is a neurodegenerative disease characterized by impairment of neuronal structures, ultimately leading to neuronal death and dysfunctions of the central nervous system. Based on previous studies, fibroblast growth factor 9 (FGF9) may provide neuroprotective functions in HD, and FGFs may enhance neuronal development and neurite outgrowth. However, whether FGF9 can provide neuronal protective functions through improvement of neuronal morphology in HD is still unclear. Here, we study the effects of FGF9 on neuronal morphology in HD and attempt to understand the related working mechanisms. Taking advantage of striatal cell lines from HD knock-in mice, we found that FGF9 increases neurite outgrowth and upregulates several structural and synaptic proteins under HD conditions. In addition, activation of nuclear factor kappa B (NF-kB) signaling by FGF9 was observed to be significant in HD cells, and blockage of NF-kB leads to suppression of these structural and synaptic proteins induced by FGF9, suggesting the involvement of NF-kB signaling in these effects of FGF9. Taken these results together, FGF9 may enhance neurite outgrowth through upregulation of NF-kB signaling, and this mechanism could serve as an important mechanism for neuroprotective functions of FGF9 in HD.

## Introduction

The basic unit of the nervous system is the neuron, which is characterized by an intricate polar morphology [1]. Proper proportioning of neuronal morphology is essential during the development of the nervous system, as it influences intracellular trafficking, synaptic development and plasticity, brain circuitry, and cognitive functions [2, 3], suggesting the importance of neuronal development under physiological conditions. Development of neuronal cells is genetically controlled by endogenous determinants as well as by exogenous signals, such as intercellular contacts, the extracellular matrix, and diffusible signals. In several neuronal diseases, abnormal development of neuronal cells or impaired neuronal morphology can be observed that lead to neuronal dysfunctions and symptoms, such as in neurodegenerative diseases [4-6]. In these diseases, the loss of neuronal structure, integrity, and connections precedes eventual death of neurons [7], suggesting that early intervention addressing neuronal morphology and synaptic functions through endogenous or exogenous factors may provide significant therapeutic benefits.

Fibroblast growth factors (FGFs) influence neuronal morphology [8-10] and are involved in the pathogenesis of many neurodegenerative diseases [11-13]. Fibroblast growth factor 9 (FGF9), a specific type of FGF, has been reported to regulate neuronal development *in vivo* [10] and also has been suggested to provide neuroprotective functions in Parkinson’s disease (PD) and Huntington’s disease (HD), two important neurodegenerative diseases [13-16]. This suggests that FGF9 may be an exogenous candidate to intervene in these diseases through regulating neuronal morphology.

Nuclear factor Kappa B (NF-kB) signaling is a ubiquitous, immediate early response that transduces extracellular signals derived from various stimuli from a variety of receptors to regulate gene expression patterns [17]. This signaling regulates a wide variety of gene expressions involved in cell growth, survival, stress responses, immune, and inflammatory processes [18]. It has also been reported that NF-kB signaling is involved in regulating cell structure, synaptic plasticity, and memory formation in the nervous system [19-21]. Furthermore, several previous studies show that NF-kB activation may be regulated by FGFs [22, 23], suggesting that FGFs might function in the morphology of neuronal cells through NF-kB signaling. In this study, we attempt to investigate the effects of FGF9 on neurite outgrowth in HD, and we also try to demonstrate the involvement of NF-kB signaling in this regulatory mechanism.

## Materials and Methods

### Cells and treatments

Institutional ethical approval was not required for this study. In this study, STHdh^Q7/Q7^(Q7) and STHdh^Q111/Q111^(Q111) striatal cells were used as the normal and HD cell models, respectively [24]. The maximum number of passages for these cell lines is 15. Q7 cells express a full-length *Huntingtin* (*HTT)* gene with 7 CAG trinucleotide repeats, and Q111 cells carry *HTT* with 111 CAG trinucleotide repeats. These cells were cultured in Dulbecco’s modified Eagle’s medium (DMEM; Gibco) with 10% (v/v) fetal bovine serum (FBS; Hyclone), L-glutamine (2 mM; Gibco), penicillin (100 U/ml; Gibco), streptomycin (100 U/ml; Gibco) and G418 (350 µg/ml; MDBio), and maintained in an incubator at 33°C under a 5% CO2 condition. For the FGF9 treatments, the cells were cultured in medium with 10% FBS for 24 hrs, and then the medium was replaced with a serum-free medium with or without FGF9 recombinant proteins (50 ng/ml; R & D Systems) for 24 or 48 hrs. To inhibit NF-kB signaling, the cells were pretreated with BAY11-7082 (1 µM; InvivoGen) 1 hr before FGF9 treatment and then cultured for 48 hrs. The cells subjected to different treatments were collected for further examination.

### Western blotting

The treated cell samples were lysed using RIPA buffer (Thermo Fisher Scientific Inc.), and crude protein was extracted using a sonicator (Qsonica) and quantitated using the Bradford assay (Pierce). These proteins were then subjected to sodium dodecyl sulfate polyacrylamide gel electrophoresis (Bio-Rad) and transferred onto PVDF membranes (Bio-Rad). These membranes were incubated with primary antibodies, including MAP2A/B (Genetex; 1:500 dilution), βIII-tubulin (Genetex; 1: 1,000 dilution), GAP-43 (Genetex; 1: 1,000 dilution), Synaptophysin (abcam; 1: 500 dilution), PSD-95 (abcam; 1:500 dilution), NF-kB/p65 (Cell signaling; 1:5,000 dilution), and γ-tubulin (Sigma; 1: 10,000 dilution) antibodies, and then secondary antibodies conjugated with peroxidase (KPL) were used to detect the primary antibodies. Protein signals were determined using a Western Lightning® Plus-ECL kit (PerkinElmer) and an imaging system (ChampGel). The quantitation of these signals was conducted using the ImageJ system (NIH).

### Immunofluorescence staining

The cells were seeded on cover glasses inside a 24-well plate and then subjected to different treatments. These treated cells were fixed with 4% paraformaldehyde for 15 mins, blocked for 1 hr and incubated with primary antibodies overnight at 4°C. The primary antibodies used in this study include βIII-tubulin (Genetex; 1: 500 dilution) and NF-kB/p65 (Cell signaling; 1:250 dilution). Afterward, secondary antibodies conjugated with fluorescent signals (Invitrogen) were incubated at room temperature for 1 hr, and then Hoechst 33342 (Sigma; 1µg/mL in PBS) was used to stain the nucleus for 15 mins at room temperature. Images were captured under a fluorescent microscope (Leica) and quantitated using the ImageJ system (NIH).

### Luciferase reporter assay

pNF-kB-Luc (Clontech), a reporter vector with the firefly luciferase gene under the control of the herpes simplex virus thymidine kinase promoter coupled with the NF-kB-binding GCCCTTAAAG sequence, was used to determine NF-kB activity. Q7 and Q111 cells were cotransfected with pNF-kB-Luc and β-galactosidase (β-gal) plasmids using Lipofectamine 2000 (Invitrogen) for 24 hrs, and then these cells were treated with or without FGF9 (50 ng/ml) for a further 24 hrs. The transfected cells were collected, and then luciferase activity was analyzed using the Dual-Luciferase^®^ Reporter Assay System (Promega). The β-gal activity was determined as an internal control to normalize luciferase activity, and the normalized results were statistically quantitated.

### Statistical Analysis

All results are presented as mean ± standard error of mean (SEM), and statistical analyses were performed using GraphPad Prism 4.02. The between-group differences were statistically analyzed using a Student’s *t*-test. Results from more than two different groups were statistically analyzed using a one-way analysis of variance (ANOVA) with Tukey’s post-hoc test. Statistical significance was shown as *p* < 0.05.

## Results

### FGF9 increases neuronal neurite outgrowth in HD cells

Previously, we demonstrated that FGF9 protects striatal cells against cell death and oxidative stress in HD [14, 15]. Since FGF9 has been reported to play a role in neuronal morphology *in vivo* [10], we were curious as to whether FGF9 would enhance neurite outgrowth and thus provide neuroprotective functions in HD cells. To examine this hypothesis, we cultured the Q7 and Q111 striatal cells from HD knock-in mice in serum-free medium with or without FGF9 for 24 and 48 hrs, and investigated the effects of FGF9 on neurite outgrowth. Based on immunofluorescent staining using a βIII-tubulin antibody, which is a neuronal marker, we found that both Q7 and Q111 cells transformed into a spindle-like shape with thinner, longer neuronal morphology after treatment with FGF9 (Figure 1A and 1B). After quantitating the total neurite outgrowth, it was found that FGF9 significantly increased the total length of the neurite outgrowth in the Q7 (Figure 1C) and Q111 (Figure 1D) cells at 24 and 48 hrs after FGF9 treatment, suggesting that FGF9 also plays a role in the development of the neuronal morphology in HD striatal cells.

**Figure 1.**
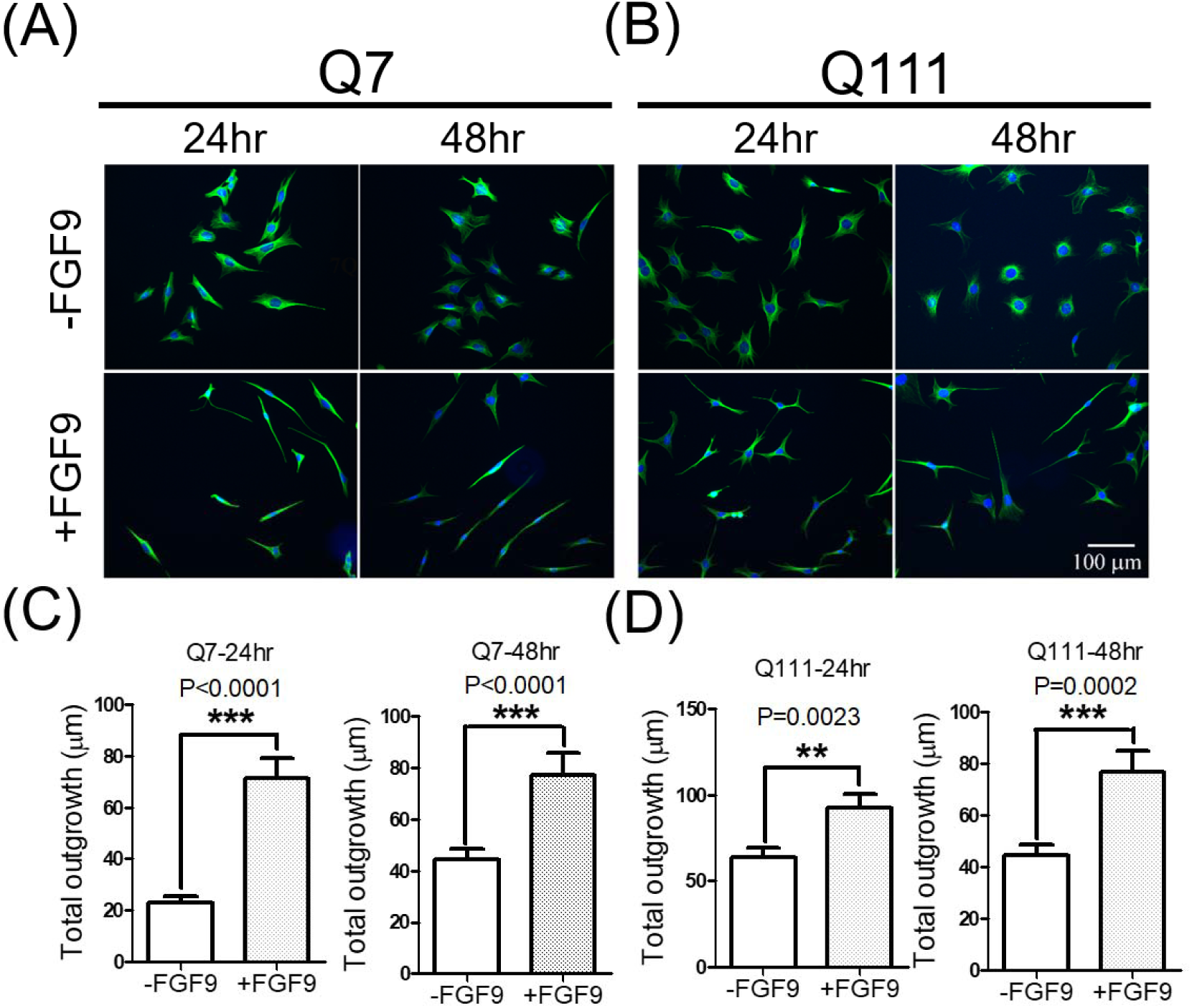
FGF9 increases neurite outgrowth in Q7 and Q111 cells. Q7 and Q111 cells were cultured with or without FGF9 for 24 and 48hrs and then fixed for immunofluorescent staining using a βIII tubulin antibody. Immunofluorescent images show neuronal morphology in Q7 **(A)** and Q111 **(B)** cells at 24 and 48 hrs after FGF9 treatment. βIII-tubulin: Green color. Hoechst 33342: Blue color. Quantitation results in the immunofluorescent images provide a comparison of neurite outgrowth in the Q7 **(C)** and Q111 **(D)** cells. ** represents *p*<0.01, *** represents *p*<0.001, N=45-103 cells from three different batches.

### FGF9 upregulates protein markers related to neuronal morphology in HD cells

The expression levels of cytoskeletal components or associated proteins, such as βIII-tubulin, microtubule associated protein (MAP2), and growth association protein 43 (GAP-43), are highly related to the development of neurite outgrowth [25, 26]. Since FGF9 increases neurite outgrowth in Q7 and Q111 cells (Figure 1), we further examined the expression profiling of these markers. Q7 and Q111 cells were treated with or without FGF9 for 48 hrs, and then subjected them to western blotting to determine the expression levels. In Q7 cells, there was no effects on the expression of βIII-tubulin, MAP2, and GAP-43 after FGF9 treatment; however, a trend in higher expression for these markers was observed (Figure 2A and 2B). In the Q111 cells, βIII-tubulin, MAP2 and GAP-43 were all upregulated by FGF9 treatment (Figure 2C and 2D), suggesting that FGF9 signaling increases the expression of these cytoskeletal components or associated proteins in HD cells.

**Figure 2.**
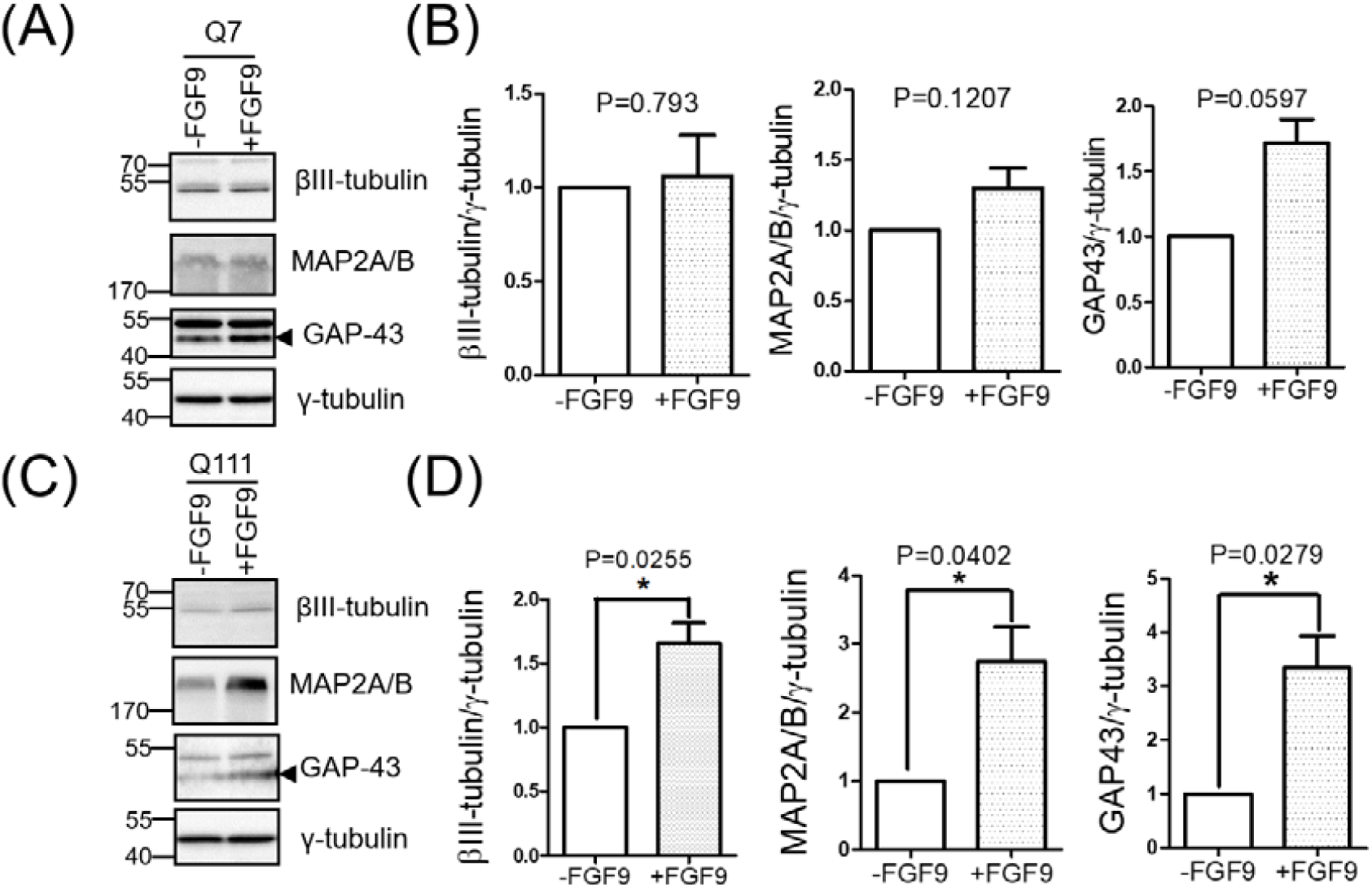
FGF9 increases expression levels of neuronal morphology-related proteins in Q7 and Q111 cells. Q7 and Q111 cells were cultured with or without FGF9 for 48hrs, and then subjected to western blotting. Western blotting was performed in Q7 **(A)** and Q111 **(C)** cells using βIII tubulin, MAP2A/B, and GAP-43 antibodies. The signal of GAP-43 is indicated by arrow heads, and γ-tubulin is used as an internal control. Quantitation results after western blotting provide a comparison of these markers in the Q7 **(B)** and Q111 **(D)** cells. * represents *p*<0.05, N= 3-4.

### FGF9 upregulates synaptic proteins in HD cells

Neurite outgrowth is highly related to neuronal functions, such as synaptic plasticity [3], and impairment of synapses is one of the neuropathological features in HD [27]. In addition, FGFs are known to promote synapse formation [28-30]. Since enhancement of neurite outgrowth was observed after FGF9 treatment of Q7 and Q111 cells, we further examined the synaptic characteristics of these treated cells through detecting two synaptic markers, synaptophysin (a presynaptic marker) and PSD-95 (a postsynaptic marker). Upon results from the western blotting, alteration of these two synaptic markers was not observed in Q7 cells (Figure 3A and 3B), whereas both synaptophysin and PSD-95 were significantly upregulated after FGF9 treatment of Q111 cells (Figure 3C and 3D). This suggests that FGF9 not only enhances neurite outgrowth, but also increases synaptic proteins.

**Figure 3.**
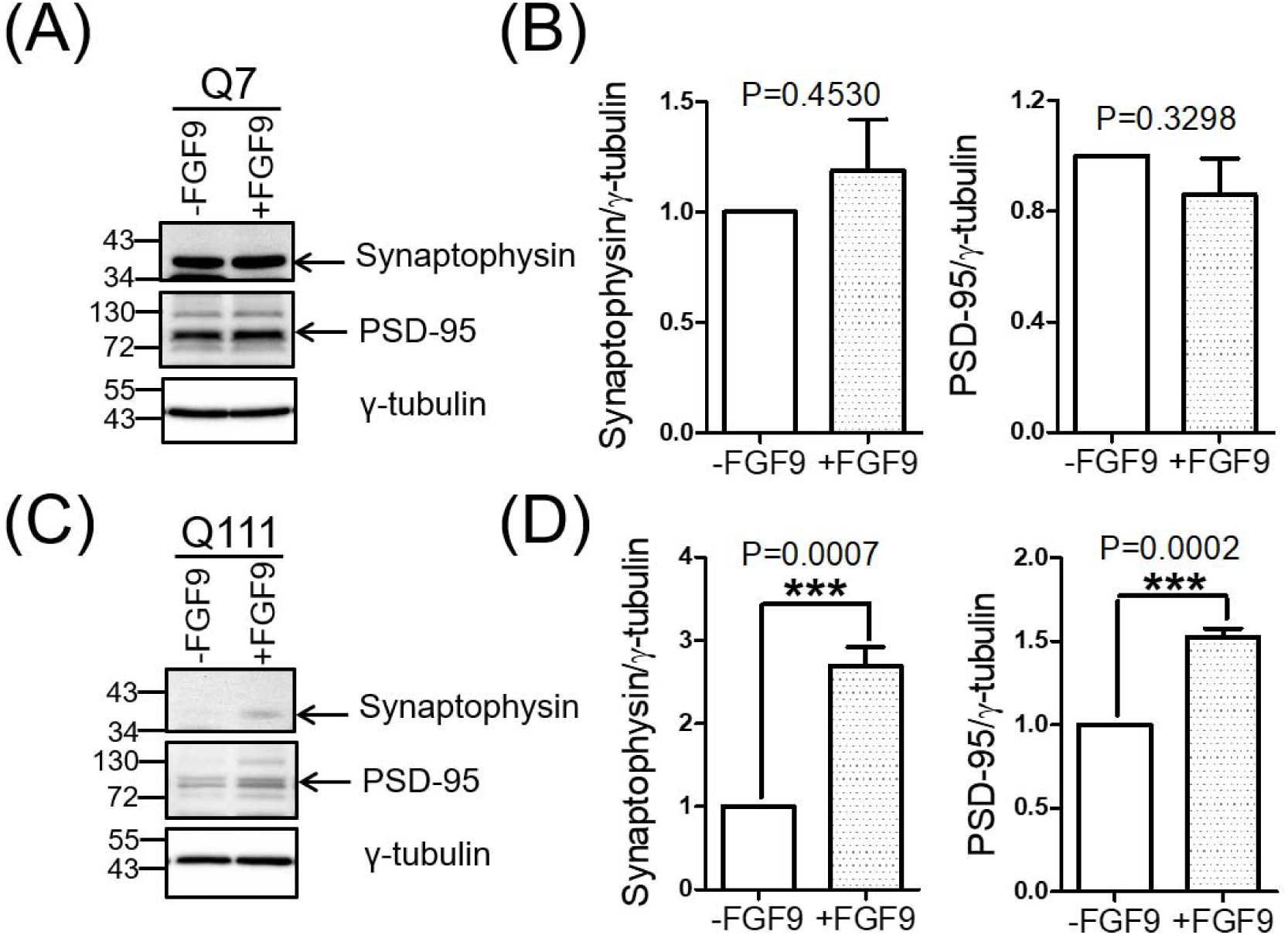
FGF9 increases expression levels of neuronal synaptic markers in Q111 cells. Q7 and Q111 cells were cultured with or without FGF9 for 48hrs and then subjected to western blotting. Western blotting was performed in the Q7 **(A)** and Q111 **(C)** cells using synaptophysin and PSD-95 antibodies. γ-tubulin was used as an internal control. Quantitation results after western blotting provide a comparison of these markers in the Q7 **(B)** and Q111 **(D)** cells. *** represents P<0.001. N= 6.

### FGF9 upregulates and activates NF-kB in HD cells

NF-kB is an important transcription factor involved in regulating several cellular processes, including neuronal morphology, synaptic development, and plasticity [18, 21, 31]. In addition, FGFs have been reported to activate NF-kB signaling in several types of cells [23, 32]. As a result, we were wondering whether FGF9 would increase neurite outgrowth through activating NF-kB signaling in HD cells. First, we attempted to determine the expression levels of NF-kB after FGF9 treatment of Q7 and Q111 cells. The two types of cells were cultured with or without FGF9 for 48 hrs, and cell samples were subjected to western blotting. The results showed that there was no change in NF-kB expression levels in the Q7 cells, but FGF9 did significantly increase the NF-kB levels in the Q111 cells (Figure 4A and 4B). Since NF-kB is a transcription factor that functions through translocating into the nucleus to activate target genes, we further attempted to examine the cellular locations of NF-kB after FGF9 treatment of HD cells. According to the results from immunostaining using an antibody against NF-kB, more NF-kB signals were observed inside the nucleus after FGF9 treatment in both the Q7 and Q111 cells (Figure 4C), and the quantitative results showed a significantly higher ratio of NF-kB signals inside the nucleus (Figure 4D), suggesting that FGF9 accelerates the translocation of NF-kB into the nucleus in both Q7 and Q111 cells. In addition, due to the fact that NF-kB functions through binding to promoter regions of target genes inside the nucleus to activate gene expression, we next determined whether FGF9 would increase NF-kB binding activity in Q7 and Q111 cells. Taking advantage of the firefly luciferase reporter assay, we transfected a reporter plasmid carrying the firefly luciferase controlled by a thymidine kinase promoter with NF-kB-binding sequences into Q7 and Q111 cells and then treated the cells with FGF9. Although we did not observe an increase in luciferase activity in the Q7 cells, FGF9 significantly increased the luciferase activity in the Q111 cells (Figure 4E). Taken these results together, FGF9 increases NF-kB expression levels, translocation into the nucleus, and promoter binding activity in HD cells.

**Figure 4.**
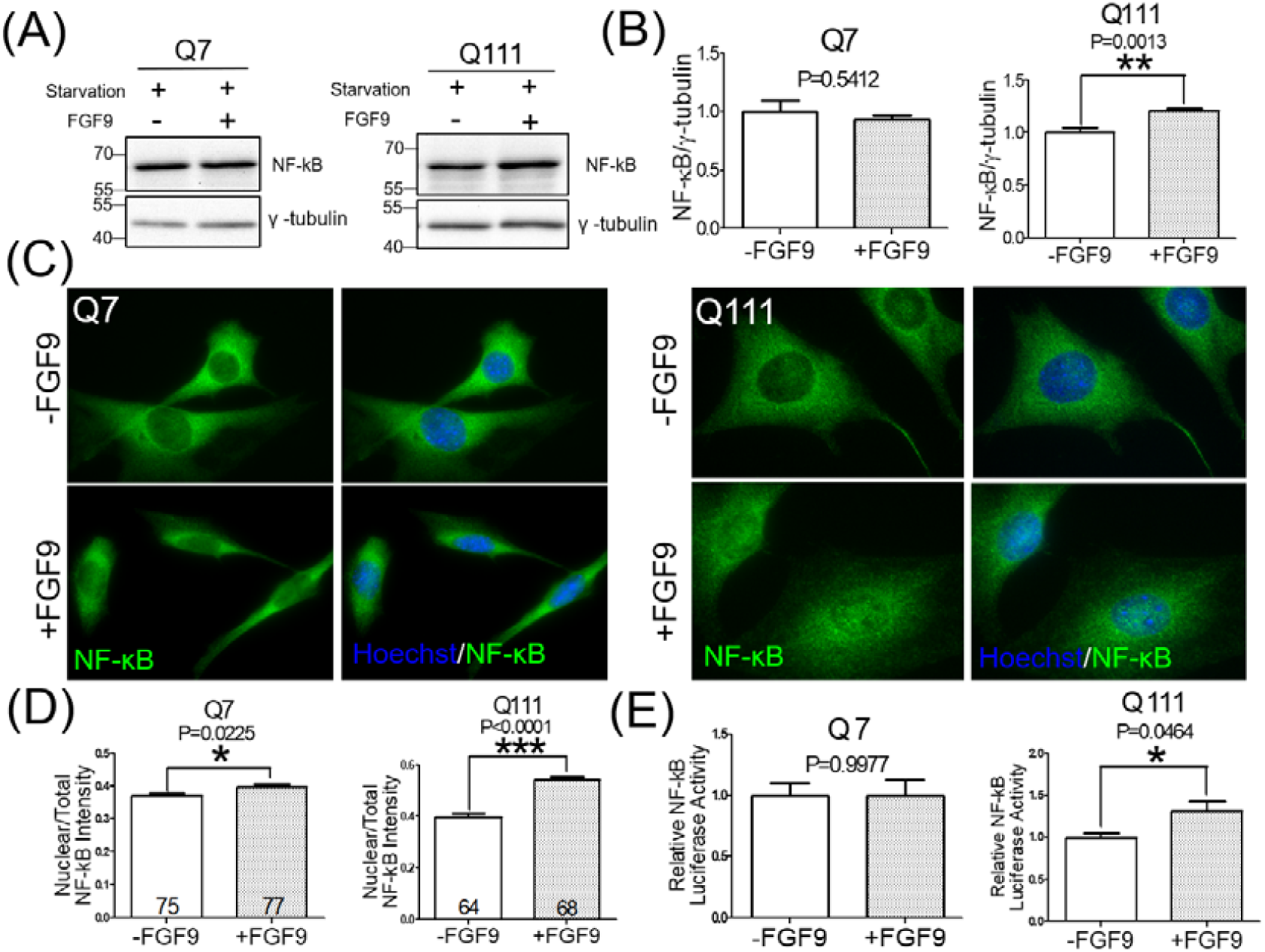
FGF9 activates NF-kB responses in Q111 cells. Q7 and Q111 cells were cultured with or without FGF9 for 48hrs and then subjected to western blotting and immunofluorescent staining. Western blotting was performed in the Q7 and Q111 cells **(A)** using a NF-kB antibody. γ-tubulin was used as an internal control. Quantitation results from (A) show a comparison of NF-kB expression in the Q7 and Q111 cells **(B)**. N= 7. Images of immunofluorescent staining using an NF-kB antibody in the Q7 and Q111 cells are shown in the Q7 and Q111 cells **(C)**. Quantitation results show the translocation of NF-kB into the nucleus in the Q7 and Q111 cells **(D)**. N= 63-77 cells from three batches. NF-kB binding activity was examined using pNF-kB-Luc plasmids, and relative NF-kB luciferase activities after FGF9 treatments in the Q7 and Q111 cells are shown in **(E)**. N=4. * represents *p*<0.05, ** represents *p*<0.01, *** represents *p*<0.001.

### FGF9 upregulates neurite outgrowth and synaptic proteins through NF-kB signaling

Based on the above results, FGF9 enhances neurite outgrowth and synaptic proteins (Figure 1-3), and it also activates NF-kB signaling in HD striatal cells (Figure 4). We therefore speculated whether FGF9 would enhance neurite outgrowth and synaptic proteins through activation of NF-kB signaling in HD cells. Using a pharmacological inhibitor, BAY11-7082, to block NF-kB activation, we examined the effects of FGF9 on the expression levels of βIII-tubulin, MAP2, and GAP-43 after interfering with the NF-kB pathway. We first confirmed that the expression level of NF-kB decreased in the Q7 and Q111 cells after treatment with of BAY11-7082 via western blotting (Supplementary Figure 1). As shown in Figure 5A and 5B, BAY11-7082 did not significantly affect the expression of βIII-tubulin in the Q7 and Q111 cells after FGF9 treatment based on the western blotting results. However, BAY11-7082 did show a trend toward decreasing the expression of βIII-tubulin induced by FGF9 in the Q111 cells (Figure 5B). Further, BAY11-7082 did not significantly decrease the expression of MAP2A/B (Figure 5C and 5D, left panel) and GAP43 (Figure 5E and 5F, left panel) in the Q7 cells, but it significantly suppressed both MAP2A/B (Figure 5C and 5D, right panel) and GAP43 (Figure 5E and 5F, right panel) induced by FGF9 in the Q111 cells. Similar to these markers related to neuronal morphology, synaptophysin and PSD-95 were also examined via western blotting. BAY11-7082 did not significantly decrease the expression of synaptophysin (Figure 6A and 6B, left panel) and PSD-95 (Figure 6C and 6D, left panel) in the Q7 cells, but it also significantly repressed both synaptophysin (Figure 6A and 6B, right panel) and PSD-95 (Figure 6C and 6D, right panel) induced by FGF9 in the Q111 cells. These findings suggest that the NF-kB pathway is important in terms of mediating FGF9 effects on neuronal morphology and synaptic proteins in HD.

**Figure 5.**
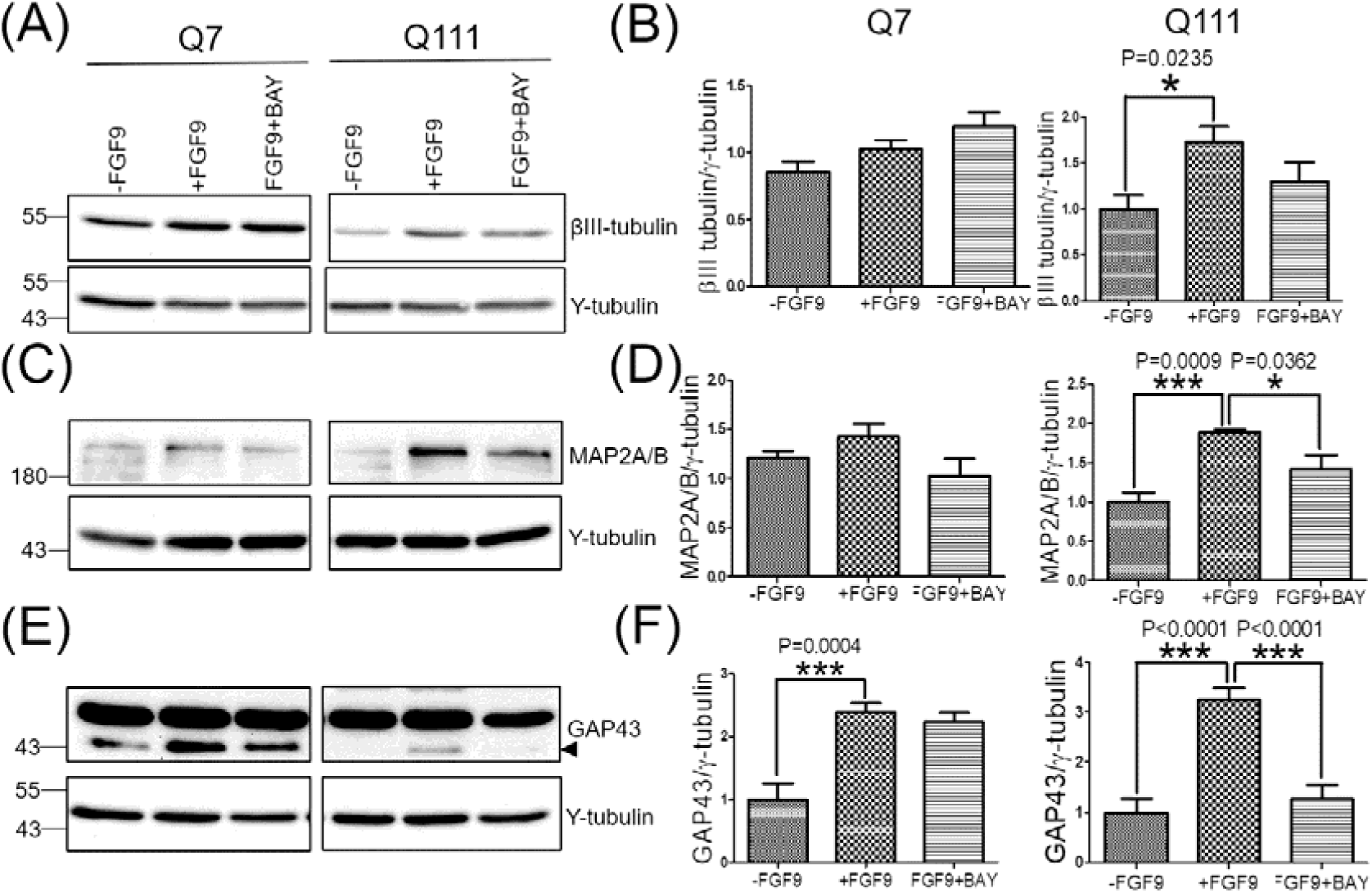
Blockage of NF-kB suppresses neuronal morphology-related proteins in Q111 cells. Q7 and Q111 cells were pretreated with BAY11-7082 for 1 hr, cultured with or without FGF9 for 48hrs, and then subjected to western blotting. Western blotting was performed in the Q7 and Q111 cells using βIII tubulin **(A)**, MAP2A/B **(C)** and GAP-43 **(E)** antibodies. The GAP-43 signal is indicated by an arrow head, and γ-tubulin is used as an internal control. Quantitation results after western blotting show a comparison of βIII tubulin **(B)**, MAP2A/B **(D)** and GAP-43 **(F)** expression in the Q7 and Q111 cells. * represents *p*<0.05, *** represents *p*<0.001. N= 4-12.

**Figure 6.**
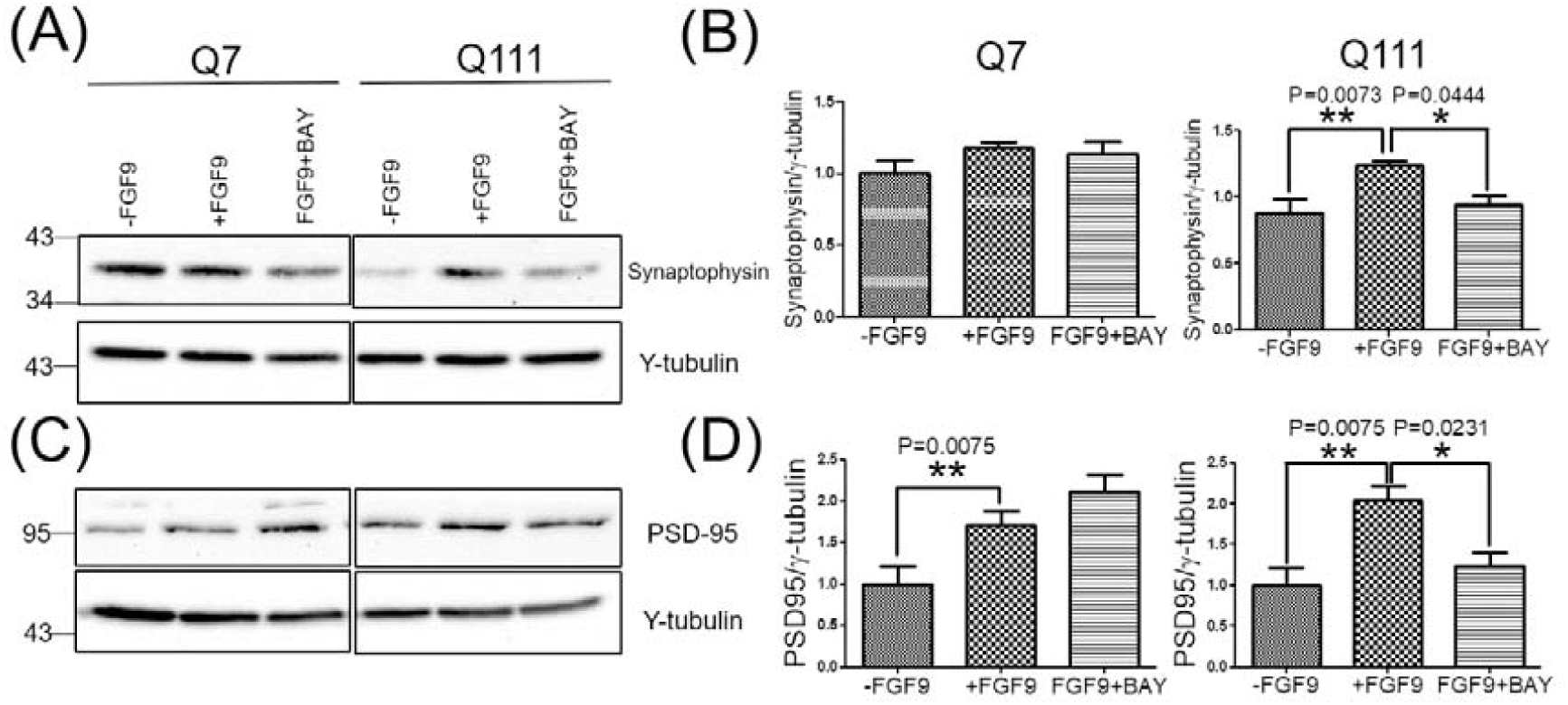
Blockage of NF-kB suppresses neuronal synaptic markers in Q111 cells. Q7 and Q111 cells were pretreated with BAY11-7082 for 1 hr, cultured with or without FGF9 for 48hrs, and then subjected to western blotting. Western blotting was performed in the Q7 and Q111 cells using synaptophysin **(A)** and PSD-95 **(C)** antibodies. γ-tubulin was used as an internal control. Quantitation results after western blotting provide a comparison of synaptophysin **(B)** and PSD-95 **(D)** expression in the Q7 and Q111 cells. * represents *p*<0.05, ** represents *p*<0.01. N= 4-7.

## Discussion

Proper proportioning of neuronal morphology is essential for nervous system development and functions, including intracellular trafficking, synaptic plasticity, brain circuitry, and ultimately, cognitive functions [2, 3]. Interestingly, impaired neuronal morphology and synaptic functions are features of many neurodegenerative diseases, including HD [4-6]. Neuronal morphological deficits precede the eventual death of neurons in these diseases [7], suggesting that early intervention addressing neuronal morphology and synaptic functions may be of significant therapeutic benefit. Previously, we reported that FGF9 has therapeutic potential by providing anti-apoptotic and anti-oxidative stress effects in HD [14, 15]. In this study, we observed that FGF9 alters neuronal morphology by increasing the total outgrowth length in Q7 and Q111 cells. We also observed that FGF9 upregulates the expression levels of specific proteins involved in regulating neuronal morphology, including βIII tubulin, MAP2A/B, GAP43, as well as synaptic proteins, including synaptophysin and PSD-95, in HD Q111 cells. In addition, we also showed that the NF-kB signaling pathway is important in terms of mediating FGF9-induced alterations in neuronal morphology and synaptic protein expression. These results together with our previous findings suggest that FGF9 provides neuroprotective functions through multiple regulatory pathways.

In the present study, we observed that FGF9 increases the total neurite outgrowth in HD cells (Figure 1). Based on previous reports, FGFs are important in regulating neuronal morphology. For example, FGF1 was reported to stimulate neurite outgrowth in PC12 cells through activating FGF receptor 1 (FGFR1) and FGF receptor 3 (FGFR3), with the latter having more potency [33]. Gain of functions for FGF9 and FGF10 has been shown in extensive dendritogenesis in the developing primary somatosensory (S1) barrel cortex [10]. FGF1 and FGF2 increase neurite outgrowth in mouse cochlear ganglion cells, and blocking FGF2 reverses this effect [34]. Furthermore, FGF2 stimulates neurite outgrowth in PC12 cells and retinal neurons, and inhibiting the FGFR1 abolishes neurite outgrowth [35]. These results are consistent with our finding indicating that FGF9 induces neurite outgrowth in HD cells. Since FGFs share several FGFRs and downstream signaling pathways, the similar neurite induction seen in these FGFs was expected. In particular, FGF2, which was reported to induce neurite outgrowth in many studies, as mentioned above, has been demonstrated to enhance neuronal survival, ameliorate neuropathological phenotypes, and ultimately increase the lifespan of R6/HD transgenic mice [36], suggesting that enhancing neuronal morphology may be one of the mechanisms for inducing protective functions in HD. However, which FGF-FGFR signaling pathway is more effective and efficient in regulating neuronal morphology is still unclear, indicating further studies are still required in this field.

Neurite specifications and neuronal development are controlled through regulating gene expressions [37, 38]. For example, FGF2 induces GAP43 protein expression and MAP2 mRNA expression in neuronal cells differentiated from bone marrow stromal cells (BMSCs) [39, 40]. FGF2 and FGF8 in combination with Sonic Hedge Hog (Shh) induce a neuron-like morphology and also stimulate the expression of neuronal markers, including neurofilament, nestin, MAP2, and βIII-tubulin, during the differentiation of human mesenchymal stem cells [41]. Consistent with previous reports, we also found that FGF9 significantly upregulates proteins involved in neuronal morphology, including βIII tubulin, MAP2A/B, and GAP43 in Q111 cells (Figure 2). Specifically, these alterations were more significantly observed in Q111 cells compared to in Q7 cells, indicating that FGF9 may more effectively improve neurite outgrowth in these disease cells and that FGF9 could be a potential target for HD therapy.

Synapses alteration is one of the neuropathological features of HD [27]. There are several studies showing the role of FGFs in synaptic development and plasticity. For example, FGF2 increases the number of synapsin I and synaptophysin positive puncta in cultured hippocampal neurons [29]. FGF22 deficiency and deletion of its receptors FGFR1 and FGFR2 resulted in limited formation of synapses in a mouse model of spinal cord injury [30]. Consistent with these reports, we found that FGF9 increases the expression of certain synaptic proteins i.e. synaptophysin and PSD-95 in Q111 cells (Figure 3), suggesting that FGF9 may efficiently enhance synaptic formations in HD cells. Similar to the structural proteins shown in Figure 2, FGF9 appears to increase expression levels in Q111 cells to a greater degree than it does in Q7 cells, suggesting the beneficial effects of FGF9 for HD disease conditions.

NF-kB is a transcription factor playing a pivotal role in regulating several cellular processes, including neuronal morphology, synaptic development and plasticity [18, 21, 31]. Previous studies have shown that this NF-kB signaling pathway is affected by FGFs in different cell types. For example, FGF1 enhances NF-kB nuclear translocation in FGFR-bearing Jurkat T cells [22], and it also increases matrix metalloproteinase-9 in mammary adenocarcinoma cells through the NF-kB pathway [32]. In addition, FGF promotes NF-kB signaling through FGFR1 in prostate cancer cells [23]. FGF2 activates NF-kB to expand the number of glial progenitor cells [42], and FGF2 inhibits TNF-mediated apoptosis in an NF-kB-dependent manner in ATCD5 cells [43]. In the current study, we found that FGF9 increases the protein expression (Figures 4A and 4B), nuclear localization (Figure 4C and 4D), and DNA-binding activity of NF-kB (Figure 4E) in Q111 cells. In addition, we also showed that BAY11-7082 significantly decreases the expression levels of morphological (Figure 5) and synaptic (Figure 6) proteins upregulated by FGF9 in Q111 cells. These results are consistent with the previous studies referenced above showing the important role of NF-kB in FGF9 signaling. However, NF-kB appears to play an ambiguous role in HD. For example, NF-kB is decreased in HD transgenic mouse and cell models, and upregulation of NF-kB increases the removal of aggregates, suggesting higher expression of NF-kB may provide beneficial effects on HD [44]. On the other hand, NF-kB is also an indicator for proinflammatory effects, and HD myeloid cells have shown an increase in NF-kB signals, leading to a cellular inflammatory response [45, 46]. We speculate that these opposite results may have been due to the use of different cell types or different stages of HD. As a result, different protective or toxic effects of NF-kB have been observed in different studies on HD. Therefore, the detailed roles of NF-kB in neuroprotection or neurotoxification in different HD cells or status should be further demonstrated.

In different studies, pharmacological inhibitors, such as BAY11-7082, are broadly used to block NF-kB in order to determine its specific roles [47-49]. Although BAY11-7082 is well-recognized as an NF-kB inhibitor, other approaches to suppress the expression of NF-kB should be used to examine FGF9-induced effects found in this study because specific drug effects should be avoided. To achieve this goal, RNA interference (RNAi), such as short interference RNA(siRNA) or short hairpin interference RNA(shRNA), can be used to inhibit the expression of NF-kB. In fact, we did try to use siRNA to knockdown the expression of NF-kB; however, the transfection efficiency of siRNAs into the Q7 and Q111 cells was very low (less than 10%). As a result, we did not observe similar blockages of FGF9-induced effects when we used siRNA against NF-kB (data not shown). Since we only used one NF-kB inhibitor, BAY11-7082, this is one drawback of this study, implying that we could not avoid the specific drug effects of BAY11-7082. To further provide more solid evidence, other pharmacological inhibitors, such as LY 294002 and Wortmannin, among others, could be used in future studies [50].

In this study, we found that FGF9-induced effects were not consistent in Q7 and Q111 cells although FGF9 increased neurite outgrowth in both cell lines (Figure 1). For example, FGF9 increases protein expression (Figures 4A and 4B), nuclear localization (Figure 4C and 4D), and DNA-binding activity of NF-kB (Figure 4E) in the Q111 cells; however, FGF9 only induced nuclear translocation of NF-kB in the Q7 cells (Figure 4C and 4D). In addition, FGF9 led to different protein expressions related to morphological and synaptic functions in the Q7 and Q111 cells (Figure 2 and 3). In several previous studies by different research groups, different responses induced by exogenous stimulations in Q7 and Q111 cells were shown [14, 51-53], suggesting these two cell lines have different regulatory networks to respond to exogenous stimulation. Since Q111 cells carry mutant HTT, which leads to disruption of gene regulation in HD models [3, 54, 55], Q111 cells may increase complementary gene regulation against HD. Oppositely, Q7 cells are wild-type cells that do not necessarily fight HD, so it would be reasonable to anticipate different responses. Also, the beneficial effects of FGF9 in terms of enhancing neurite outgrowth through NF-kB signaling were more obviously observed in the Q111 cells, but a significant enhancement in neurite outgrowth was also recorded in the Q7 cells (Figure 1). Because we only examined specific representative markers related to neuronal morphology, other structural or synaptic markers or morphology-related signals may be increased significantly after treatment with FGF9 in Q7 cells. This might explain the conflicting results in the Q7 cells. In future studies, it would be interesting to further investigate the differences in the detailed mechanisms affected by FGF9 in these two types of cells.

In summary, FGF9 not only stimulates neurite outgrowth but also increases synaptic proteins in striatal cell models of HD. Most importantly, FGF9 offers these functions through NF-κB signaling under the HD condition, which is one of the important mechanisms for neuroprotective functions of FGF9 in HD. Based on our series of studies, we provide several important regulatory mechanisms of FGF9 for neuroprotection in HD and hope to accelerate the applications of FGF9 in this devastating disease.

## Lists of abbreviations

(HD): Huntington’s disease
(FGF9): Fibroblast growth factor 9
(NF-kB): Nuclear factor kappa B
(PD)\: Parkinson’s disease
(Q7): STHdh^Q7/Q7^
(Q111): STHdh^Q111/Q111^
(MAP2): Microtubule associated protein
(GAP-43): Growth association protein 43
(FGFR): FGF receptor
(Shh): Sonic Hedge Hog

## Ethics approval and consent to participate

Not applicable

## Consent for publication

Not applicable

## Availability of data and materials

The datasets and materials used and/or analysed during the current study are available from the corresponding author upon reasonable request.

## Authors and Contributors

IOY, HMC, PHC, and CYC handled the cellular studies and molecular analysis and analyzed the data; SJT, JIC, CCW, BMH, HSS, CMC and SHY designed the experiments and oversaw the progress of the study. IOY, HMC and SHY drafted the paper. All authors read and approved the final manuscript.

## Acknowledgements

Not applicable

## Figure legends

**Supplementary Figure 1.**
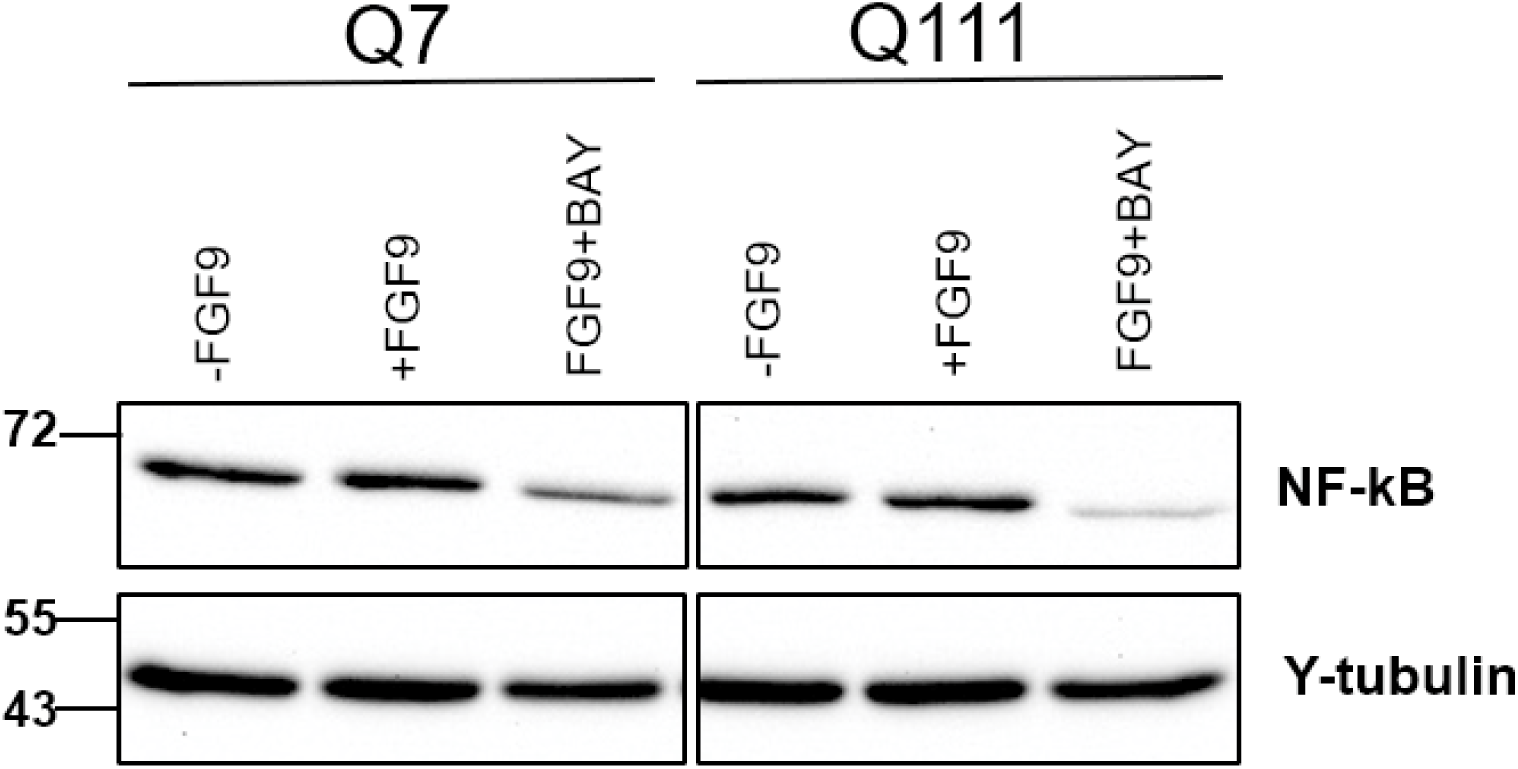
BAY11-7082 suppresses the expression of NF-kB in Q7 and Q111 cells with the treatment of FGF9. Q7 and Q111 cells were pretreated with BAY11-7082 for 1 hr, cultured with or without FGF9 for 48hrs, and then subjected to western blotting using NF-kB and γ-tubulin antibodies. γ-tubulin was used as an internal control.

